# Avian uropygial secretions as a novel bioinsecticide for mosquito vector control

**DOI:** 10.64898/2026.10.13.757564

**Authors:** A. Marzal, Y. Sato, S. Magallanes, A. Díaz, J. Muriel, M. Ferraguti, J. Martínez-de la Puente, E. Frontera, C Núñez, Y. Tsuda, M. Inumaru, L. García-Longoria

## Abstract

Mosquito-borne diseases remain a major global health burden, yet current vector-control programmes are threatened by the rapid emergence of resistance to conventional insecticides. The urgent need for new, effective, and environmentally sustainable insecticidal agents has promoted growing interest in nature-derived compounds. Birds produce uropygial secretions with well-documented antimicrobial properties, but their potential to directly kill mosquito vectors has never been experimentally tested. Here, we assess whether uropygial secretions from house sparrows act as naturally derived insecticides against five mosquito species responsible for the global transmission of human pathogens (*Anopheles stephensi*, *Culex pipiens, Aedes aegypti*, *Aedes albopictus*, and *Culiseta longiareolata*). We show that uropygial secretions induce high mortality (65–94%) across all five mosquito species in a dose- and time-dependent manner. These findings provide the first experimental evidence that avian preen secretions contain previously unrecognized bioactive compounds capable of rapidly killing mosquitoes, revealing a novel biological resource with direct relevance for public health.

## INTRODUCTION

More than 80% of the global population is at risk of contracting vector-borne diseases (1), which impose substantial burdens on public health systems and global economies (2). Among these, mosquito-borne diseases represent the greatest impact, accounting for approximately 350 million cases and over one million deaths annually (3). In recent years, incidence rates of diseases such as malaria and dengue have risen sharply (3), driven in part by the widespread emergence of resistance to WHO-recommended insecticides. This resistance compromises the effectiveness of key interventions, including insecticide-treated nets and indoor residual spraying (4).

Insecticide resistance has now been documented across multiple mosquito taxa of major medical importance. For instance, many populations of *Culex pipiens*, the primary vector of pathogens such as West Nile virus, Rift Valley fever virus, and filarial parasites, exhibit resistance to organochlorine insecticides (5). Similarly, resistance to pyrethroids, organophosphates, and carbamate insecticides has been documented in Asia for *Culiseta longiareolata*, a species of medical importance due to its role in transmitting arboviruses (6). Resistance has also spread rapidly among malaria vectors in endemic regions, potentially enhancing vector competence and transmission capacity of *Plasmodium* parasites (7, 8). Notably, *Anopheles stephensi*, a primary malaria vector in South Asia and the Arabian Peninsula, and an emerging invasive species in Africa, has already developed resistance to insecticides used in indoor residual spraying and long-lasting insecticidal nets (9; 10). Furthermore, resistance or elevated tolerance to insecticides has been reported worldwide in *Aedes aegypti* and *Aedes albopictus*, the principal vectors for arboviruses causing chikungunya, dengue, Zika, and yellow fever (9; 11; 12). Collectively, these trends underscore the urgent need for innovative tools and sustainable vector-control strategies.

The uropygial gland (also known as *preen gland*) is an exocrine gland located at the base of the tail in most bird species. It produces a lipid-rich secretion used primarily for plumage maintenance and known to possess antimicrobial properties (13; 14). Over the past decade, several studies have suggested an additional defensive role of these uropygial secretions against malaria infection. For instance, Magallanes et al. (15) found that uninfected house sparrows (*Passer domesticus*) had larger uropygial glands and exhibited higher antimicrobial activity in their secretions compared to infected individuals. Marzal et al. (16) reported similar patterns in sparrow populations from both Peru and Spain, with uninfected birds showing both larger uropygial glands and uropygial secretions with stronger antibacterial activity than infected conspecifics. In a broader comparative analysis across 37 bird species from temperate (Spain) and tropical (Peru) regions, Magallanes et al. (17) likewise reported that infected individuals generally had smaller uropygial glands than uninfected birds. These findings suggest that the uropygial gland secretions may constitute a defence against malaria infection, potentially influencing the interaction between birds and their haemosporidian vectors through insecticidal (18; 14) or repellent effects (19; but see 20). However, direct evidence of insecticidal activity against mosquito vectors remains lacking.

In this study, we experimentally test the insecticidal potential of uropygial secretions from a widespread passerine, the house sparrows, by assessing their effects on five mosquito species spanning four different genera: *An. stephensi*, *Cx. pipiens*, *Ae. aegypti*, *Ae. albopictus* and *Cu. longiareolata.* These mosquitoes include both native and invasive species and are considered major contributors to the global transmission of human vector-borne pathogens. Specifically, quantify mosquito mortality across a range of secretion concentrations. Demonstrating insecticidal activity in uropygial secretions would identify a previously overlooked source of bioactive compounds and challenge the prevailing view that these secretions function primarily in antimicrobial defence. From a public health perspective, such findings could lay the foundation for developing innovative, biologically inspired tools to complement existing vector-control strategies (21).

## RESULTS

Uropygial gland secretions exhibited a strong insecticidal effect across all five mosquito species tested. Mortality varied significantly with both secretion concentration and exposure time, with higher concentrations and longer exposure consistently resulting in increased mortality compared to controls (Table 2 and Figure 1).

**Table 1.** Final concentrations of uropygial gland secretion (UGS) solutions used in CDC Bottle Bioassays.

| **Solution** |  |
| --- | --- |
| 0.5× | 50 µl UGS + 50 µl ddH<sub>2</sub>O |
| 0.25× | 25 µl UGS + 75 µl ddH<sub>2</sub>O |
| 0.125× | 12.5 µl UGS + 85.5 µl ddH<sub>2</sub>O |
| Control | 100 µl ddH<sub>2</sub>O |

**Table 2.**
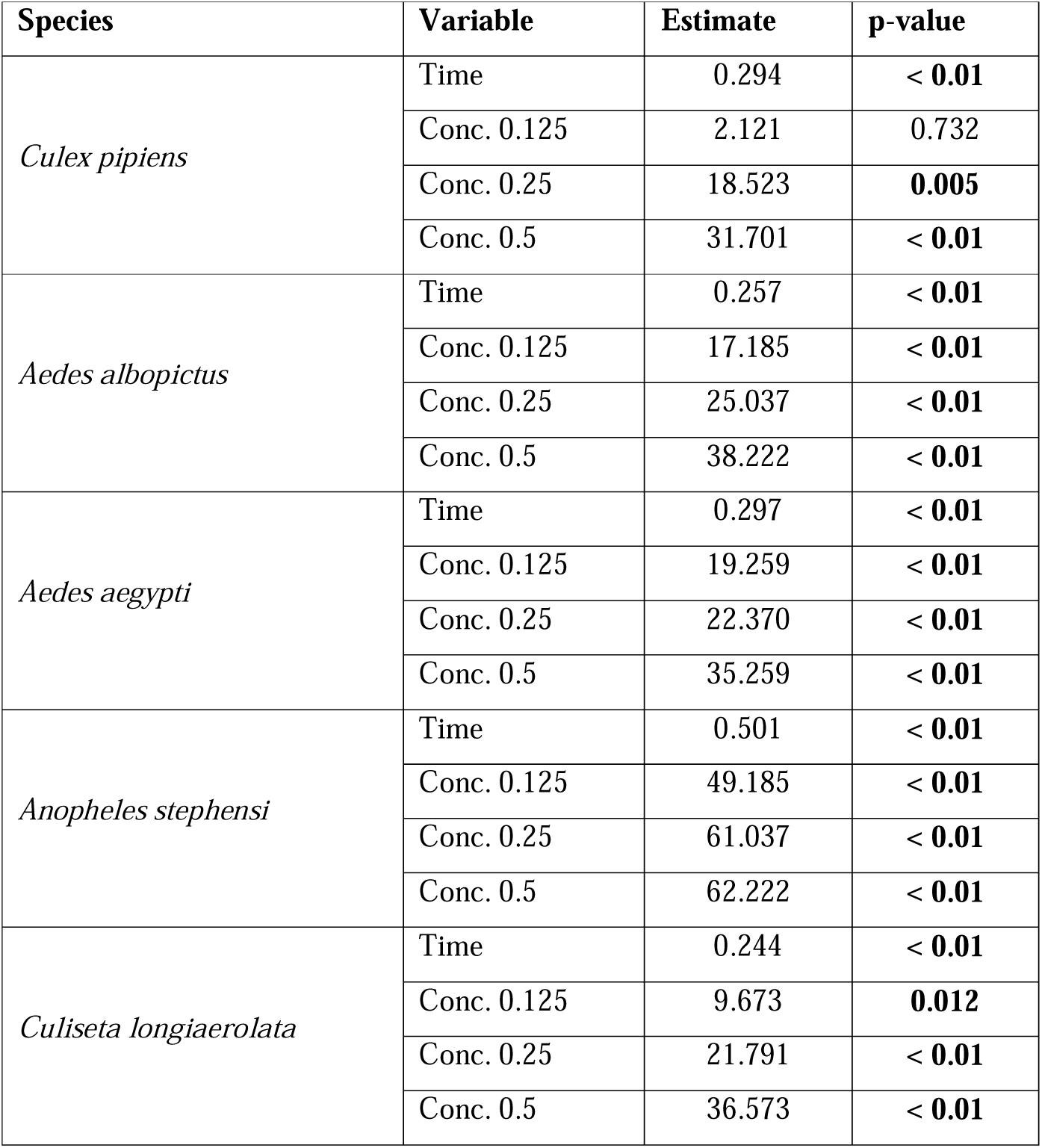
Statistical results of mosquito knockdown/mortality across species. The table presents the effects of time and different concentrations on mosquito knockdown/mortality for five species: *Culex pipiens*, *Aedes albopictus*, *Aedes aegypti*, *Anopheles stephensi*, and *Culiseta longiaerolata*. The columns display the analyzed variable (time or concentration), the estimated effect, and the corresponding p-value.

**Figure 1.**
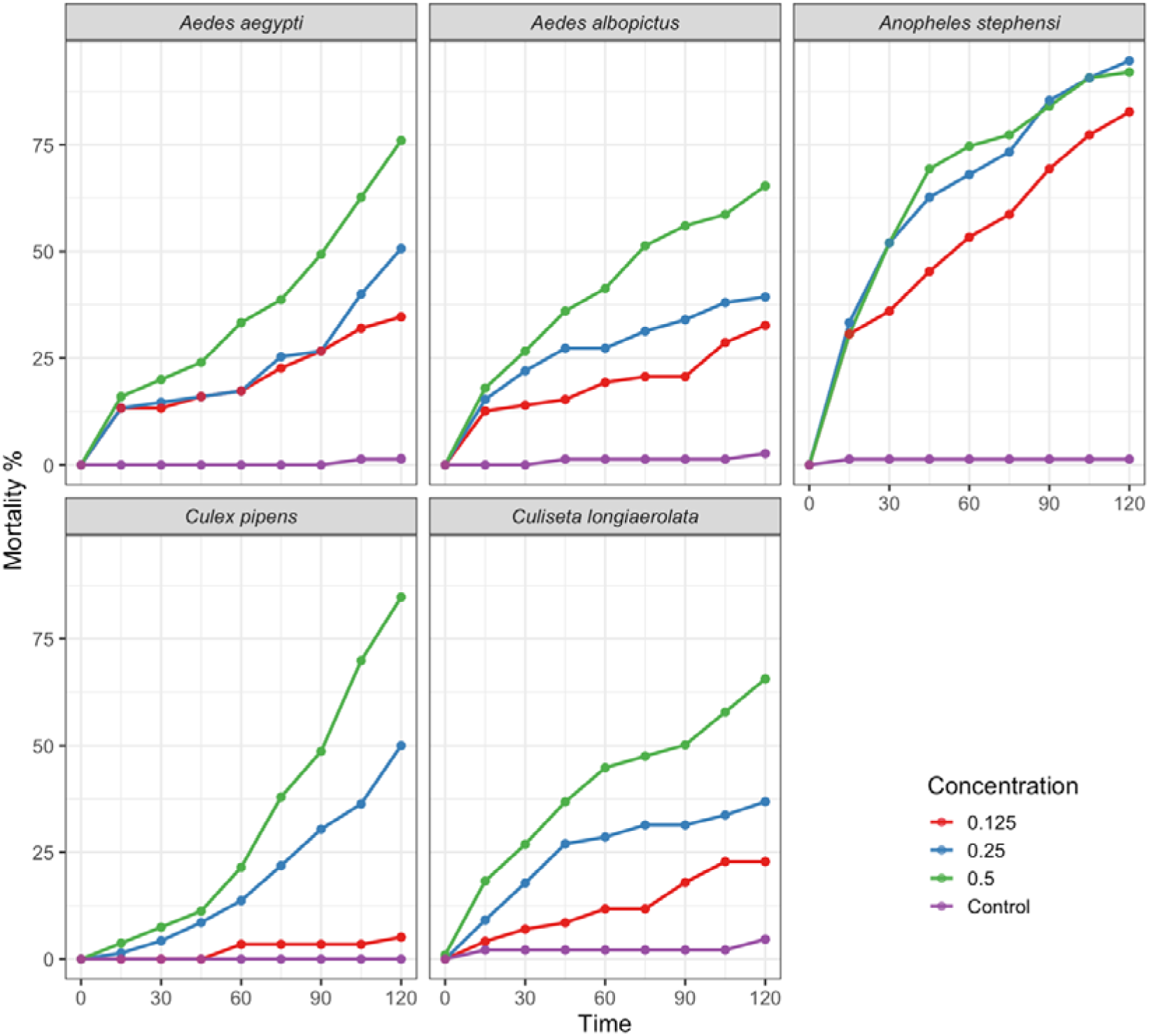
Mortality percentages of the five mosquito species over time (min) when exposed to three concentrations of uropygial secretion from house sparrows (*Passer domesticus*) and the control. Each line represents a different uropigial gland concentration. Mortality rates were recorded at multiple time points.

In *Ae. aegypti*, the model explained 80.22% of the variation in mortality. Exposure to 0.5×, 0.25×, and 0.125× concentrations of uropygial secretions significantly increased mortality relative to controls, with mortality rising progressively over exposure time (Table 2, Figure 1A). Similar patterns were observed in *Ae. albopictus* (R² = 83.73%), where both exposure time and secretion concentration were strong predictors of mosquito mortality (Table 2, Figure 1B).

In *An. stephensi*, the model explained 84.53% of the variation in mortality, revealing strong effects of both preen oil concentration and exposure time during bioassays on mosquito mortality. Mortality reached maximum values of 92%, 94%, and 83% at 0.5×, 0.25×, and 0.125× concentrations, respectively (Table 2, Figure 1C).

For *Cx. pipiens*, 66.88% of the variation in mortality was explained by the model. Mortality reached maximum values of 92%, 94%, and 83% at 0.5×, 0.25×, and 0.125× concentrations, respectively (Table 2, Figure 1D).

In *Cs. longiaerolata*, the model explained 84.24% of the variability in mortality. Similar to the patterns observed in the other species, mortality increased with both rising secretion concentration and longer exposure (Table 2, Figure 1E).

Overall, mosquito mortality varied significantly among species after 120 min of exposure to uropygial secretion across all tested concentrations (χ² = 24, df = 4, p = 7.99 × 10). Post-hoc analyses revealed marked interspecific differences in susceptibility to preen oil, with *An. stephensi* being particularly sensitive, even at low concentrations, while *Cs. longiareolata* consistently exhibited the lowest mortality rate (Table 3, Figure 2).

**Table 3:**
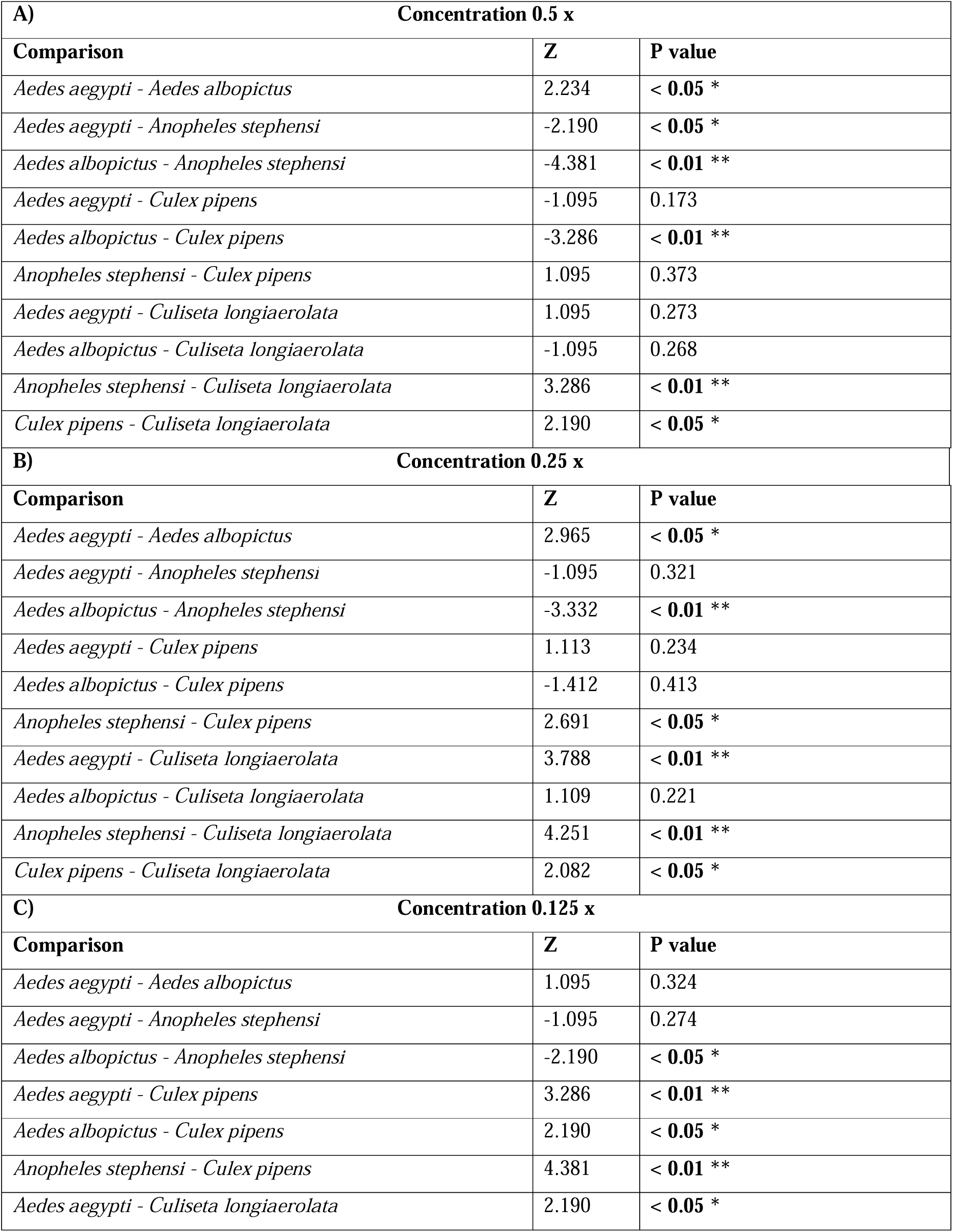
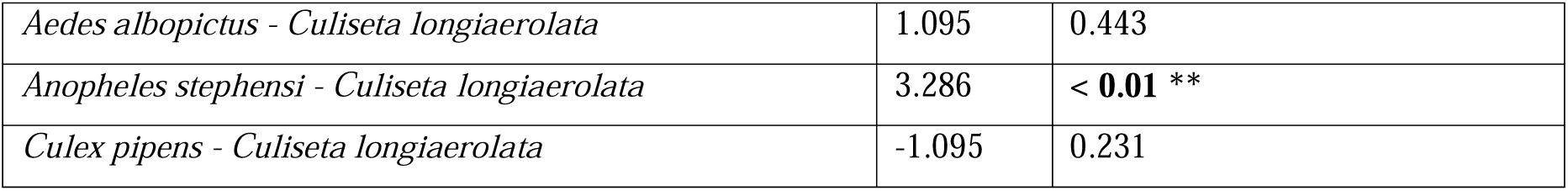
Pairwise post hoc comparisons of mosquito species knockdown/mortality after 120 minutes of exposure to three concentrations (0.5×, 0.25×, and 0.125×) (A;B;C, respectively) of uropygial gland secretion. The table shows Z-scores and corresponding p-values for each species comparison at each concentration. Asterisks indicate the level of statistical significance (*p* < 0.05: *, *p* < 0.01: **).

**Figure 2.**
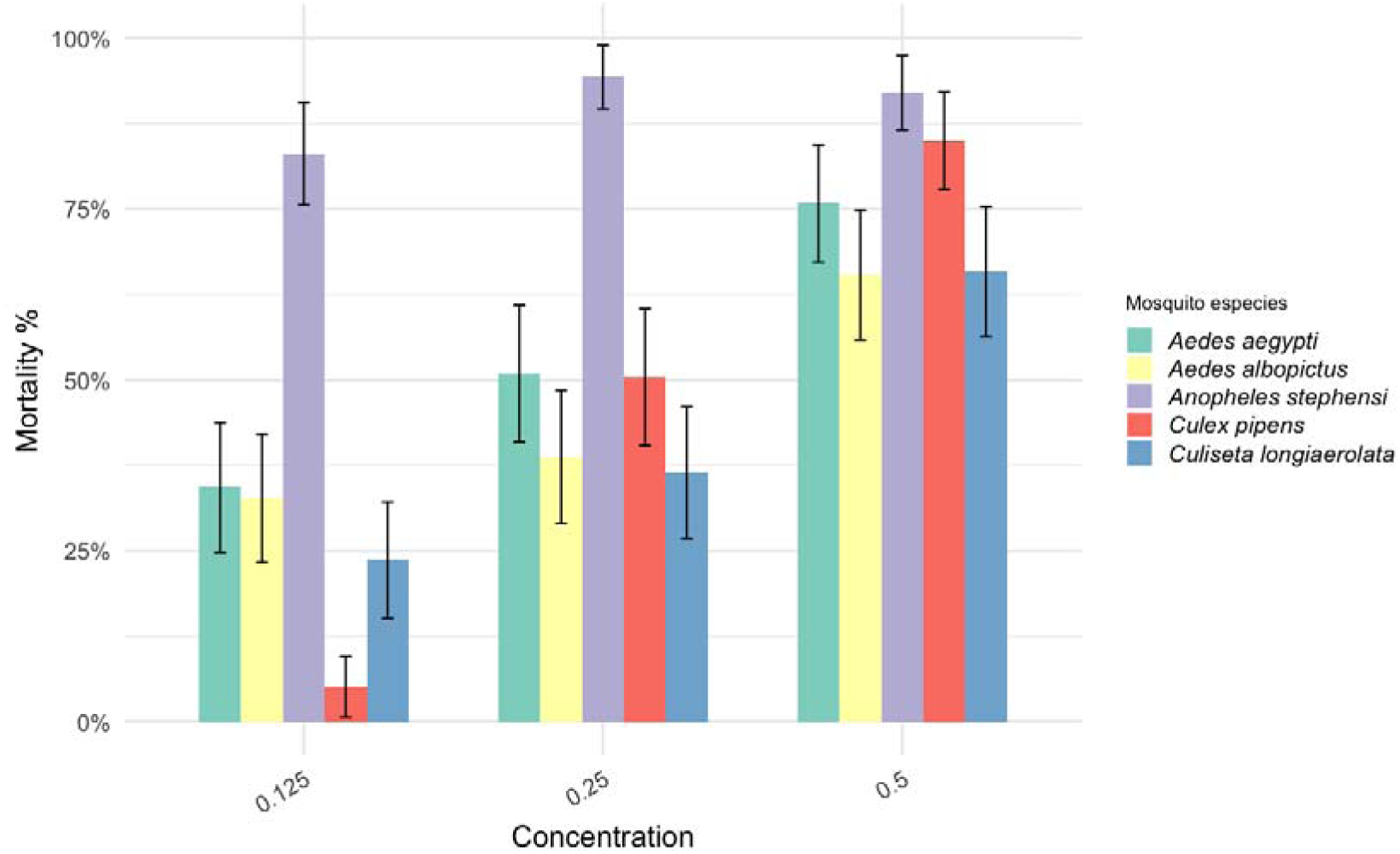
Mortality rates of mosquito species after 120 minutes at three different concentrations. The bars represent the mean mortality rate (%), and the error bars indicate the standard error (SE). Different colours represent different species of mosquitoes. For statistical results, please refer to Table S1.

## DISCUSSION

The global rise of insecticide resistance represents a major public health challenge, undermining the effectiveness of vector-control programs and threatening progress in disease prevention (4). In response, the World Health Organization has recently emphasized the urgent need to discover and deploy new insecticides to sustain advances against vector-borne diseases such as malaria (3). In this context, our study demonstrates that uropygial gland secretions from house sparrows induce substantial mortality in five mosquito species that are major vectors of severe or fatal diseases in humans and animals. Importantly, these findings challenge the long-standing assumption that uropygial secretions primarily serve antimicrobial or maintenance functions, instead revealing that they can contain potent vector-toxic compounds. These findings highlight the potential role of gland-derived compounds as promising bioinsecticides and open new opportunities for developing biologically inspired, next-generation vector-control tools that could complement conventional approaches.

The observed dose- and time-dependent effects suggest that the toxic activity of uropygial secretions scales with both concentration and exposure duration. This pattern is consistent with known mechanisms of action in other insecticides, where toxicity depends on cumulative exposure (22; 23). Although the precise mode of action remains to be determined, several hypotheses can be proposed. For example, uropygial secretions may impair ectoparasites by coating their surfaces or occluding spiracles, leading to suffocation (24). Alternatively, specific chemical constituents of avian secretion, such 2-tridecanone and hexadecanoic acid, have demonstrated insecticidal activity (25; 26) and have been detected in the preen secretions of numerous bird species (see review in 27). Further investigations should focus on isolating and characterizing these active components and experimentally validating their insecticidal effects.

Notably, susceptibility to uropygial secretion varied among mosquito species. *Anopheles stephensi* consistently showed the highest sensitivity, with substantial mortality even at the concentration (0.125×). In contrast, *Cx. pipiens* and *Cs. longiareolata* exhibited greater resilience, particularly at intermediate and low doses. *Aedes aegypti* and *Ae. albopictus* displayed moderate sensitivity, with mortality rising mainly at higher concentrations but remaining below than observed in *An. stephensi*. These species-specific differences are consistent with broader evidence that mosquito susceptibility to insecticides is influenced by differences in physiology, metabolism, and ecology (28). For example, habitat preferences and exposure to environmental stressors differ between *Culex* and *Anopheles*, potentially shaping responses to bioactive secretions (29). *Anopheles stephensi* typically breeds in relatively clean, stagnant waters (30), and often shows greater physiological sensitivity to environmental stressors (31), which may explain its vulnerability. In contrast, *Cx. pipiens* thrives in polluted, organically rich waters (32; 33) and may encounter a broader suite of chemical stressors (34; 35), fostering stronger detoxification mechanisms and greater resilience to uropygial compounds.

Additional mechanisms contributing to reduced susceptibility may include enhanced metabolic resistance through the overexpression or upregulation of detoxification enzymes that degrade insecticides such as esterases and cytochrome P450 monooxygenases, which are linked to widespread insecticide resistance in *Culex* mosquitoes (36). Also, target-site insensitivity comprises mutations that reduce insecticide binding or efficacy at the site of action, such as voltage-gated sodium channel mutations associated with pyrethroid resistance and organochlorine resistance observed across numerous *Culex*, *Aedes*, and *Anopheles* populations worldwide (36; 37; 38). Moreover, structural differences in the cuticle, such as increased thickness or reduced permeability that limit insecticide penetration (e.g., thicker femoral cuticles in resistant vs susceptible *Cx. pipiens*), have been reported (39). Finally, behavioural differences in resting patterns can influence exposure, with *Anopheles* species tending to maintain longer contact with treated surfaces, whereas *Culex* species often exhibit avoidance behaviour (40).

In conclusion, our findings identify uropygial gland secretions as a promising source of novel, naturally derived insecticidal compounds. These secretions represent a previously overlooked biological resource with potential applications in vector control. Their development could contribute to alternative or complementary tools within integrated vector-management programmes, reducing reliance on synthetic insecticides. More broadly, this study highlights the potential of leveraging naturally occurring compounds to diversify vector-control strategies and address the growing challenge of insecticide resistance. Further research should aim to isolate active compounds, elucidate their mechanisms of action, and evaluate their safety, efficacy, and scalability under field conditions.

## MATERIAL AND METHODS

### Collection of uropygial gland secretions

Between February and March 2018, we captured 134 wild adult house sparrows using mist nets in Badajoz, Spain. Immediately after capture, we collected the entire available uropygial secretion from the gland papilla of each individual following the protocol described by Magallanes et al. (15). Briefly, the uropygial gland and surrounding area were first cleaned with a cotton swab soaked in ethanol to minimize contamination risk. After the ethanol evaporated, the papilla was gently pressed with a finger to expel the secretion, which was then collected into a microcapillary tube until the papilla was empty. The secretions from all individuals were pooled, transferred into a sterile Eppendorf vial, and kept at approximately 4°C in a portable icebox during the fieldwork. They were subsequently stored at -20°C in the laboratory within 4 hours until use in Centers for Disease Control and Prevention (CDC) bottle bioassays.

### Mosquito larvae collection and mosquito rearing

Mosquito larvae from *Cx. pipiens* and *Cs. longiaerolata* were collected using standard dippers and nets in September – October 2018 from small freshwater ponds in Badajoz (38°39’N, 7°13’W) and from marshlands in Huelva Province (37°17′N, 6°53′W), Spain, respectively. Larvae from *An. stephensi*, *Ae. aegypti* and *Ae. albopictus* were obtained in March 2019 from outbred colonies established at the Department of Medical Entomology, National Institute of Infectious Diseases (NIID), Japan.

Larvae were transported to the corresponding laboratories: *Cx. pipiens* to the University of Extremadura (UEx), *Cs. longiareolata* to the Doñana Biological Station (EBD), and *An. stephensi*, *Ae. aegypti*, and *Ae. albopictus* to NIID, and reared under controlled conditions. Thet were supplied with shrimp food (GmbH & Co. KG, D-53501) and allowed to develop in plastic trays filled with water inside climatic chambers set at 28±1°C, 60–65% relative humidity, and a 12:12h light:dark cycle.

After emergence, adult mosquitoes maintained according to standard protocol for mosquito colony maintenance described in Kauffmann et al. (41). Briefly, they were kept were kept in insect-rearing cages (BugDorm-43030F) at temperatures between 25-30°C and relative humidity of 65–70%, under a 12:12h light:dark photoperiod, with *ad libitum* access to a 5% sugar solution until the experiments. Adult mosquitoes used in the assays were 10–14 days old. Bioassays were conducted at UEx for *Cx. pipiens* (September 2018), at EBD for *Cs. longiareolata* (October 2018), and at NIID for *Ae. aegypti, Ae. albopictus,* and *An. stephensi* (March 2019).

### CDC Bottle Bioassay

The insecticidal effect of uropygial secretion was evaluated following the CDC Bottle Bioassay protocol (42). Although originally developed to detect insecticide resistance in mosquitoes by applying a controlled dose of insecticide coated on the interior surface of a bottle, this assay can also determine whether a particular substance at a given concentration induces mosquito mortality within a specified time frame (43; 44).

In total, 2293 mosquitoes were tested for susceptibility to uropygial secretion (476 *Cx. pipiens*, 411 *Cu. longiareolata*, 465 *Ae. aegypti*, 489 *Ae. albopictus*, and 452 *An. stephensi*). We conducted 25 bottle bioassays (five per species) to quantify mortality following exposure to uropygial secretion. Each bioassay included four replicate bottles: three containing different concentrations of the uropygial gland secretion (0.5×, 0.25×, and 0.125×) and one untreated control (Table 1). The bioassay protocol was as follows: first, 100 µl of either ddH O (control) or uropygial gland secretion solution at the specified concentration (treatments) was added to each bottle using a 200 µl micropipette with disposable tips (Table 1). Second, the bottle caps were securely tightened, then gently swirled, inverted, and rotated to ensure even coating on all sides of the interior surface. Third, immediately after coating bottles with uropygial solutions or ddH O, 20 – 25 adult mosquitoes were aspirated from the insect-rearing cages and gently introduced into each bottle. Control bottles were loaded first, followed by the treatment bottles. All bioassays were performed with bottles placed on their sides to ensure consistency across trials.

### Mortality assessment and data validity

Time zero was recorded upon mosquito introduction. Mosquito activity was monitored at 15-min intervals over a 120-min exposure period, and the numbers of live and knocked-down individuals were recorded at each interval. Mosquitoes were scored as dead if unable to stand on their legs or were immobile, sliding along the curvature of the bottle. The mortality of mosquitoes in control groups from all bioassays was below 3%, thus validating the data in the experiments (42).

### Statistical analyses

To assess the effects of time and concentration on mosquito mortality, linear models (LM) were fitted separately for each mosquito species. The response variable was mortality (%), while time (continuous) and concentration (categorical, with "Control" as the reference level) were included as predictor variables. Model assumptions, including normality and homoscedasticity of residuals, were tested using residual diagnostic analyses performed with the *ggfortify* (45) and *performance* (46) packages. The significance of model coefficients was evaluated using the *t-statistics* and corresponding *p-values*. Model fit was assessed using the adjusted R-squared and F-statistic. To compare mosquito mortality across species, we conducted chi-squared tests on mortality rates after 120 minutes for each concentration. Pairwise comparisons between species were further examined using post hoc Bonferroni-adjusted tests. All analyses were conducted using the *stats* package in R (version 4.4.0; 47) and figures were generated with ggplot2 package (48).

## ACKNOWLEDGEMENTS

This paper is dedicated to the memory of our dear co-author Dr. Yoshio Tsuda, who sadly passed away in 2024.

## AUTHOR CONTRIBUTIONS

AM, LG-L and YS conceived the idea of the study. All authors collected the samples and performed the experiments. AM and LGL analyzed the data, created the figures and tables, and wrote the first draft of the manuscript. All authors read, contributed to, and approved the final version of the manuscript.

## DECLARATION OF GENERATIVE AI IN SCIENTIFIC WRITING

During the preparation of this work the authors used Grammarly (app.grammarly.com) in order to enhance the language quality. After using this tool, the authors reviewed and edited the content as needed and took full responsibility for the content of the published article.

## DATA STATEMENT

All data presented in this manuscript are available and can be obtained from the corresponding author upon reasonable request.

## Abstract

### Background

Malaria control relies heavily on interventions targeting mosquito vectors, but the rapid emergence of insecticide resistance threatens the effectiveness of existing approaches. Identifying novel, effective, and environmentally sustainable insecticides is therefore a priority. Birds produce uropygial secretions with well-documented antimicrobial properties, but their potential insecticidal activity against mosquitoes, including malaria vectors, remains unexplored.

### Methods

We evaluated the insecticidal activity of uropygial secretions from wild house sparrows (*Passer domesticus*) against the malaria vector *Anopheles stephensi* and four additional mosquito species of medical importance (*Aedes aegypti, Aedes albopictus, Culex pipiens*, and *Culiseta longiareolata*). A total of 2,293 adult mosquitoes were tested using CDC bottle bioassays at three secretion concentrations (0.5×, 0.25×, and 0.125×) and an untreated control. Mosquito mortality was monitored for 120 min.

### Results

Uropygial secretions caused substantial mosquito knockdown/mortality, with effects increasing with concentration and exposure time. Susceptibility differed significantly among species after 120 min. Notably, *An. stephensi* was particularly susceptible, reaching knockdown/mortality of 92%, 94%, and 83% at 0.5×, 0.25×, and 0.125× concentrations, respectively.

### Conclusions

Our findings provide the first experimental evidence that avian uropygial secretions possess direct insecticidal activity against mosquitoes and identify them as a previously overlooked source of naturally derived insecticidal compounds. The high susceptibility of *An. stephensi* highlights their potential relevance for developing novel bioinsecticides for malaria vector control.

## INTRODUCTION

More than 80% of the global population is at risk of contracting vector-borne diseases [1], which impose substantial burdens on public health systems and global economies [2]. Among these, mosquito-borne diseases represent the greatest impact, accounting for approximately 350 million cases and over one million deaths annually [3]. Malaria remains one of the most important mosquito-borne diseases worldwide, with vector control representing a cornerstone of malaria prevention and elimination strategies [4]. However, in recent years, the incidence of malaria and other mosquito-borne diseases, such as dengue, has increased substantially, while the widespread emergence of resistance to WHO-recommended insecticides threatens the effectiveness of current vector-control interventions [3, 4]. In particular, insecticide resistance compromises key malaria-control tools, including insecticide-treated nets and indoor residual spraying [5].

Insecticide resistance has now been documented across multiple mosquito taxa of major medical importance. For instance, resistance has spread rapidly among malaria vectors in endemic regions, potentially enhancing vector competence and transmission capacity of *Plasmodium* parasites [6, 7]. Notably, *Anopheles stephensi*, a primary malaria vector in South Asia and the Arabian Peninsula, and an emerging invasive species in Africa, has already developed resistance to insecticides used in indoor residual spraying and long-lasting insecticidal nets [8]. Also, many populations of *Culex pipiens*, the primary vector of pathogens such as West Nile virus, Rift Valley fever virus, and filarial parasites, exhibit resistance to organochlorine insecticides [9]. Similarly, resistance to pyrethroids, organophosphates, and carbamate insecticides has been documented in Asia for *Culiseta longiareolata*, a species of medical importance due to its role in transmitting arboviruses [10]. Furthermore, resistance or elevated tolerance to insecticides has been reported worldwide in *Aedes aegypti* and *Aedes albopictus*, the principal vectors for arboviruses causing chikungunya, dengue, Zika, and yellow fever [11]. Collectively, these trends underscore the urgent need for innovative tools and sustainable vector-control strategies.

The uropygial gland (also known as *preen gland*) is an exocrine gland located at the base of the tail in most bird species. It produces a lipid-rich secretion used primarily for plumage maintenance and is known to possess antimicrobial properties [12]. Over the past decade, several studies have suggested an additional defensive role of these uropygial secretions against malaria infection. For instance, Magallanes et al. [13] found that uninfected house sparrows (*Passer domesticus*) had larger uropygial glands and exhibited higher antimicrobial activity in their secretions compared to infected individuals. Marzal et al. [14] reported similar patterns in sparrow populations from both Peru and Spain, with uninfected birds showing both larger uropygial glands and uropygial secretions with stronger antibacterial activity than infected conspecifics. In a broader comparative analysis across 37 bird species from temperate (Spain) and tropical (Peru) regions, Magallanes et al. [15] likewise reported that infected individuals generally had smaller uropygial glands than uninfected birds. Together, these findings suggest that uropygial gland secretions may contribute to defence against avian malaria not only through their antimicrobial properties, but also by influencing interactions between birds and their haemosporidian vectors through repellent or insecticidal effects [16] (but see [17]). However, whether these secretions exert direct insecticidal effects on mosquito vectors has never been experimentally tested.

In this study, we experimentally test the insecticidal potential of uropygial secretions from house sparrows against five mosquito species spanning four genera: *An. stephensi*, *Cx. pipiens*, *Ae. aegypti*, *Ae. albopictus* and *Cs. longiareolata*. These species include important vectors of malaria and other human pathogens. Using CDC bottle bioassays, we quantify mosquito knockdown/mortality across different secretion concentrations and exposure times. We hypothesize that uropygial secretions exert direct insecticidal effects and that susceptibility varies among mosquito species. Demonstrating such activity would identify a previously overlooked source of naturally derived insecticidal compounds, expanding the recognized defensive functions of avian uropygial secretions beyond their well-established antimicrobial properties. From a public health perspective, these compounds could provide a basis for developing biologically inspired tools to complement existing strategies for malaria and broader mosquito-vector control [18].

## METHODS

### Collection of uropygial gland secretions

Between February and March 2018, we captured 134 wild adult house sparrows using mist nets in Badajoz, Spain. Immediately after capture, we collected the entire available uropygial secretion from the gland papilla of each individual following the protocol described by Magallanes et al. [13]. Briefly, the uropygial gland and surrounding area were first cleaned with a cotton swab soaked in ethanol to minimize contamination risk. After the ethanol evaporated, the papilla was gently pressed with a finger to expel the secretion, which was then collected into a microcapillary tube until the papilla was empty. The secretions from all individuals were pooled, transferred into a sterile Eppendorf vial, and kept at approximately 4°C in a portable icebox during the fieldwork. They were subsequently stored at -20°C in the laboratory within 4 hours until use in Centers for Disease Control and Prevention (CDC) bottle bioassays.

### Mosquito larvae collection and mosquito rearing

Larvae were transported to the corresponding laboratories: *Cx. pipiens* to the University of Extremadura (UEx), *Cs. longiareolata* to the Doñana Biological Station (EBD), and *An. stephensi*, *Ae. aegypti*, and *Ae. albopictus* to NIID, and reared under controlled conditions. They were supplied with shrimp food (GmbH & Co. KG, D-53501) and allowed to develop in plastic trays filled with water inside climatic chambers set at 28±1°C, 60–65% relative humidity, and a 12:12h light:dark cycle.

After emergence, adult mosquitoes were maintained according to standard protocol for mosquito colony maintenance described in Kauffmann et al. [19]. Briefly, they were kept were kept in insect-rearing cages (BugDorm-43030F) at temperatures between 25-30°C and relative humidity of 65–70%, under a 12:12h light:dark photoperiod, with *ad libitum* access to a 5% sugar solution until the experiments. Adult mosquitoes used in the assays were 10–14 days old. Bioassays were conducted at UEx for *Cx. pipiens* (September 2018), at EBD for *Cs. longiareolata* (October 2018), and at NIID for *Ae. aegypti, Ae. albopictus,* and *An. stephensi* (March 2019).

### CDC Bottle Bioassay

The insecticidal effect of uropygial secretion was evaluated following the CDC Bottle Bioassay protocol [20]. Although originally developed to detect insecticide resistance in mosquitoes by applying a controlled dose of insecticide coated on the interior surface of a bottle, this assay can also determine whether a particular substance at a given concentration induces mosquito mortality within a specified time frame [21].

In total, 2293 mosquitoes were tested for susceptibility to uropygial secretion (476 *Cx. pipiens*, 411 *Cu. longiareolata*, 465 *Ae. aegypti*, 489 *Ae. albopictus*, and 452 *An. stephensi*). We conducted 25 bottle bioassays (five per species) to quantify mortality following exposure to uropygial secretion. Each bioassay included four replicate bottles: three containing different concentrations of the uropygial gland secretion (0.5×, 0.25×, and 0.125×) and one untreated control (Table 1). The bioassay protocol was as follows: first, 100 µl of either ddH₂O (control) or uropygial gland secretion solution at the specified concentration (treatments) was added to each bottle using a 200 µl micropipette with disposable tips (Table 1). Second, the bottle caps were securely tightened, then gently swirled, inverted, and rotated to ensure even coating on all sides of the interior surface. Third, immediately after coating bottles with uropygial solutions or ddH₂O, 20 – 25 adult mosquitoes were aspirated from the insect-rearing cages and gently introduced into each bottle. Control bottles were loaded first, followed by the treatment bottles. All bioassays were performed with bottles placed on their sides to ensure consistency across trials.

### Knockdown / Mortality assessment and data validity

Time zero was recorded upon mosquito introduction. Mosquito activity was monitored at 15-min intervals over a 120-min exposure period, and the numbers of active and knocked-down/immobile individuals were recorded at each interval. Mosquitoes were classified as knocked down when they were unable to stand on their legs or remained immobile and slid along the curvature of the bottle. Because no post-exposure recovery period was included, this endpoint does not distinguish irreversible mortality from temporary knockdown. Accordingly, the response measured in the bioassays is hereafter referred to as knockdown/mortality. Knockdown/mortality in control groups was below 3% in all bioassays, supporting the validity of the experiments [20].

### Statistical analyses

To assess the effects of time and concentration on mosquito knockdown/mortality, linear models (LM) were fitted separately for each mosquito species. The response variable was knockdown/mortality (%), while time (continuous) and concentration (categorical, with "Control" as the reference level) were included as predictor variables. Model assumptions, including normality and homoscedasticity of residuals, were tested using residual diagnostic analyses performed with the *ggfortify* [22] and *performance* [23] packages. The significance of model coefficients was evaluated using the *t-statistics* and corresponding *p-values*. Model fit was assessed using the adjusted R-squared and F-statistic. To compare mosquito knockdown/mortality across species, we conducted chi-squared tests on knockdown/mortality rates after 120 minutes for each concentration. Pairwise comparisons between species were further examined using post hoc Bonferroni-adjusted tests. All analyses were conducted using the *stats* package in R (version 4.4.0) [24] and figures were generated with ggplot2 package [25].

## RESULTS

Uropygial gland secretions exhibited a strong insecticidal effect across all five mosquito species tested. Knockdown/mortality varied significantly with both secretion concentration and exposure time, with higher concentrations and longer exposure consistently resulting in increased knockdown/mortality compared to controls (Table 2 and Figure 1).

In *An. stephensi*, the model explained 84.53% of the variation in knockdown/mortality, revealing strong effects of both preen oil concentration and exposure time during bioassays on mosquito mortality. Knockdown/mortality reached maximum values of 92%, 94%, and 83% at 0.5×, 0.25×, and 0.125× concentrations, respectively (Table 2, Figure 1).

In *Ae. aegypti*, the model explained 80.22% of the variation in knockdown/mortality. Exposure to 0.5×, 0.25×, and 0.125× concentrations of uropygial secretions significantly increased knockdown/mortality relative to controls, with knockdown/mortality rising progressively over exposure time (Table 2, Figure 1). Similar patterns were observed in *Ae. albopictus* (R² = 83.73%), where both exposure time and secretion concentration were strong predictors of mosquito knockdown/mortality (Table 2, Figure 1).

For *Cx. pipiens*, 66.88% of the variation in knockdown/mortality was explained by the model. knockdown/mortality reached maximum values of 92%, 94%, and 83% at 0.5×, 0.25×, and 0.125× concentrations, respectively (Table 2, Figure 1).

In *Cs. longiaerolata*, the model explained 84.24% of the variability in knockdown/mortality. Similar to the patterns observed in the other species, knockdown/mortality increased with both rising secretion concentration and longer exposure (Table 2, Figure 1E).

Overall, mosquito knockdown/mortality varied significantly among species after 120 min of exposure to uropygial secretions across all tested concentrations (χ² = 24, df = 4, *p* = 7.99 × 10⁻). Post-hoc analyses revealed marked interspecific differences in susceptibility to uropygial secretions. Among the five species, *An. stephensi* showed particularly high susceptibility, including substantial knockdown/mortality at the lowest tested concentration, whereas *Cs. longiareolata* consistently exhibited the lowest response (Table 3, Figure 2).

## DISCUSSION

The global rise of insecticide resistance represents a major public health challenge, undermining the effectiveness of vector-control programs and threatening progress in disease prevention [4]. In response, the World Health Organization has recently emphasized the urgent need to discover and deploy new insecticides to sustain advances against vector-borne diseases such as malaria [5]. In this context, our study demonstrates that uropygial gland secretions from house sparrows induce substantial mortality in five mosquito species that are major vectors of severe or fatal diseases in humans and animals. Importantly, these findings challenge the long-standing assumption that uropygial secretions primarily serve antimicrobial or maintenance functions, instead revealing that they can contain potent vector-toxic compounds. These findings highlight the potential role of gland-derived compounds as promising bioinsecticides and open new opportunities for developing biologically inspired, next-generation vector-control tools that could complement conventional approaches.

The observed dose- and time-dependent effects suggest that the toxic activity of uropygial secretions scales with both concentration and exposure duration. This pattern is consistent with known mechanisms of action in other insecticides, where toxicity depends on cumulative exposure [26, 27]. Although the precise mode of action remains to be determined, several hypotheses can be proposed. For example, uropygial secretions may impair ectoparasites by coating their surfaces or occluding spiracles, leading to suffocation [28]. Alternatively, specific chemical constituents of avian secretion, such 2-tridecanone and hexadecanoic acid, have demonstrated insecticidal activity [29, 30] and have been detected in the preen secretions of numerous bird species (see review in [17]). Further investigations should focus on isolating and characterizing these active components and experimentally validating their insecticidal effects.

Notably, susceptibility to uropygial secretion varied among mosquito species. *Anopheles stephensi* consistently showed the highest sensitivity, with substantial mortality even at the concentration (0.125×). In contrast, *Cx. pipiens* and *Cs. longiareolata* exhibited greater resilience, particularly at intermediate and low doses. *Aedes aegypti* and *Ae. albopictus* displayed moderate sensitivity, with knockdown/mortality rising mainly at higher concentrations but remaining below than observed in *An. stephensi*. These species-specific differences are consistent with broader evidence that mosquito susceptibility to insecticides is influenced by differences in physiology, metabolism, and ecology [31]. For example, habitat preferences and exposure to environmental stressors differ between *Culex* and *Anopheles*, potentially shaping responses to bioactive secretions [32]. *Anopheles stephensi* typically breeds in relatively clean, stagnant waters [33], and often shows greater physiological sensitivity to environmental stressors [34], which may explain its vulnerability. In contrast, *Cx. pipiens* thrives in polluted, organically rich waters [35] and may encounter a broader suite of chemical stressors [36], fostering stronger detoxification mechanisms and greater resilience to uropygial compounds.

Additional mechanisms contributing to reduced susceptibility may include enhanced metabolic resistance through the overexpression or upregulation of detoxification enzymes that degrade insecticides such as esterases and cytochrome P450 monooxygenases, which are linked to widespread insecticide resistance in *Culex* mosquitoes [37]. Also, target-site insensitivity comprises mutations that reduce insecticide binding or efficacy at the site of action, such as voltage-gated sodium channel mutations associated with pyrethroid resistance and organochlorine resistance observed across numerous *Culex*, *Aedes*, and *Anopheles* populations worldwide [37-39]. Moreover, structural differences in the cuticle, such as increased thickness or reduced permeability that limit insecticide penetration (e.g., thicker femoral cuticles in resistant vs susceptible *Cx. pipiens*), have been reported [40]. Finally, behavioural differences in resting patterns can influence exposure, with *Anopheles* species tending to maintain longer contact with treated surfaces, whereas *Culex* species often exhibit avoidance behaviour [41].

Several limitations should be considered when interpreting these findings. The active compound(s) responsible for mosquito mortality remain unidentified, and their mechanisms of action are unknown. Moreover, CDC bottle bioassays demonstrate activity under controlled laboratory conditions but do not establish efficacy under field conditions. Potential toxicity to non-target organisms and the environmental persistence and scalability of candidate compounds also remain to be evaluated. Finally, differences in the origin and rearing history of the mosquito species tested may have contributed to interspecific variation in susceptibility.

In conclusion, our findings identify uropygial gland secretions as a promising source of novel, naturally derived insecticidal compounds. These secretions represent a previously overlooked biological resource with potential applications in vector control. Their pronounced activity against *An. stephensi* is particularly relevant given the increasing importance of this invasive malaria vector and the growing challenge of insecticide resistance. Their development could contribute to alternative or complementary tools within integrated vector-management programmes, reducing reliance on synthetic insecticides. More broadly, this study highlights the potential of leveraging naturally occurring compounds to diversify vector-control strategies and address the growing challenge of insecticide resistance. Isolation and characterization of the active compounds, followed by assessment of their mode of action, safety, formulation and efficacy under field conditions, will determine whether these naturally occurring molecules can contribute to future integrated malaria vector-control strategies.

## DATA STATEMENT

The data supporting this study are available from the corresponding author upon reasonable request.

## Legends for figures

Figure 1. **Mosquito knockdown/mortality over time following exposure to uropygial secretions.** Knockdown/mortality (%) of five mosquito species during 120 min of exposure to three concentrations (0.5×, 0.25×, and 0.125×) of uropygial gland secretion from house sparrows (*Passer domesticus*) or to the control (ddH₂O). Responses were recorded at 15-min intervals.

Figure 2. **Mosquito knockdown/mortality after 120 min of exposure to uropygial secretions**. Mean knockdown/mortality (%) of five mosquito species after exposure to three concentrations (0.5×, 0.25×, and 0.125×) of uropygial gland secretion from house sparrows (*Passer domesticus*). Error bars represent standard errors (SE). Statistical comparisons among species are reported in Table 3.

## REFERENCES

1. Franklinos LHV, Jones KE, Redding DW, Abubakar I. The effect of global change on mosquito-borne disease. Lancet Infect Dis. 2019 Sep;19(9):e302–e312. doi: 10.1016/S1473-3099(19)30161-6. Epub 2019 Jun 18. PMID: 31227327.

2. Teeluck, M., Adegboye, O., Karl, S., Iyaloo, D. P., & McBryde, E. (2025). Understanding the Effect of a Changing Climate on the Re Emergence of Mosquito Borne Diseases in Vulnerable Small Island Nations: A Systematic Review. Zoonoses and Public Health.

3. WHO guidelines for malaria, 13 August 2025. Geneva: World Health Organization; 2025. 10.2471/B09514. Licence: CC BY-NC-SA 3.0 IGO.

4. Van den Berg, H., da Silva Bezerra, H.S., Al-Eryani, S., et al. Recent trends in global insecticide use for disease vector control and potential implications for resistance management. Sci Rep 11, 23867 (2021). 10.1038/s41598-021-03367-9

5. Abbasi, E., & Daliri, S. (2024). Knockdown resistance (kdr) associated organochlorine resistance in mosquito-borne diseases (Culex quinquefasciatus): systematic study of reviews and meta-analysis. PLOS Neglected Tropical Diseases, 18(8), e0011991.

6. Soltanbeiglu, S., Bavani, M. M., Vahedi, M., & Chavshin, A. R. (2022, September). Susceptibility of Culiseta longiareolata to insecticides in West Azerbaijan province, northwestern Iran. In Veterinary Research Forum (Vol. 13, No. 3, p. 435).

7. Suh, P. F., Elanga-Ndille, E., Tchouakui, M., Sandeu, M. M., Tagne, D., Wondji, C., & Ndo, C. (2023). Impact of insecticide resistance on malaria vector competence: a literature review. Malaria Journal, 22(1), 19.

8. Oruni, A., Tchouakui, M., Tagne, C. S. D., Hearn, J., Kayondo, J., & Wondji, C. S. (2024). Temporal evolution of insecticide resistance and bionomics in Anopheles funestus, a key malaria vector in Uganda. Scientific Reports, 14(1), 32027.

9. Teshome, A., Erko, B., Golassa, L. et al. Resistance of Anopheles stephensi to selected insecticides used for indoor residual spraying and long-lasting insecticidal nets in Ethiopia. Malar J 22, 218 (2023). 10.1186/s12936-023-04649-5

10. Taylor, R., Messenger, L. A., Abeku, T. A., Clarke, S. E., Yadav, R. S., & Lines, J. (2024). Invasive Anopheles stephensi in Africa: insights from Asia. Trends in parasitology.

11. Zulfa, R., Lo, W. C., Cheng, P. C., Martini, M., & Chuang, T. W. (2022). Updating the insecticide resistance status of Aedes aegypti and Aedes albopictus in Asia: a systematic review and meta-analysis. Tropical medicine and infectious disease, 7(10), 306.

12. Asgarian, T. S., Vatandoost, H., Hanafi-Bojd, A. A., & Nikpoor, F. (2023). Worldwide status of insecticide resistance of Aedes aegypti and Ae. albopictus, vectors of arboviruses of Chikungunya, Dengue, Zika and Yellow Fever. Journal of Arthropod-Borne Diseases, 17(1), 1.

13. Moreno Rueda, G., 2017. Preen oil and bird fitness: a critical review of the evidence. Biological Reviews, 92(4), pp.2131–2143

14. Mora Rubio, C., García Longoria, L., & Marzal, A. (2025). Preening for protection: a systematic review of the antimicrobial properties of uropygial secretions. Journal of Avian Biology, 2025(2), e03425.

15. Magallanes, S., Møller, A. P., García-Longoria, L., de Lope, F., & Marzal, A. (2016). Volume and antimicrobial activity of secretions of the uropygial gland are correlated with malaria infection in house sparrows. Parasites & Vectors, 9(1), 232.

16. Marzal, A., Møller, A. P., Espinoza, K., Morales, S., Luján Vega, C., Cárdenas Callirgos, J. M., … & Magallanes, S. (2018). Variation in malaria infection and immune defence in invasive and endemic house sparrows. Animal Conservation, 21(6), 505–514.

17. Magallanes, S., MøLLER, A. P., Lujan-Vega, C., Fong, E., Vecco, D., Flores-Saavedra, W., … & Marzal, A. (2021). Exploring the adjustment to parasite pressure hypothesis: differences in uropygial gland volume and haemosporidian infection in palearctic and neotropical birds. Current Zoology, 67(2), 147–156.

18. Marzal, A., Magallanes, S., & Garcia-Longoria, L. (2022). Stimuli followed by avian malaria vectors in host-seeking behaviour. Biology, 11(5), 726.

19. Martínez-de la Puente, J., Díez-Fernández, A., Soriguer, R. C., Rambozzi, L., Peano, A., Meneguz, P. G., & Figuerola, J. (2020). Are malaria-infected birds more attractive to mosquito vectors?. Ardeola, 68(1), 205–218.

20. Díez Fernández, A., Martínez de la Puente, J., Gangoso, L., Ferraguti, M., Soriguer, R., & Figuerola, J. (2020). House sparrow uropygial gland secretions do not attract ornithophilic nor mammophilic mosquitoes. Medical and Veterinary Entomology, 34(2), 225–228.

21. Cai, P., & Dimopoulos, G. (2024). Microbial biopesticides: A one health perspective on benefits and risks. One Health, 100962.

22. Sovi A, Adoha CJ, Yovogan B, Cross CL, Dee DP, Konkon AK, Sidick A, Accrombessi M, Ahouandjinou MJ, Ossè R, Dangbénon E, Towakinou L, Agbangla C, Padonou GG, Churcher TS, Ngufor C, Cook J, Protopopoff N, Akogbéto MC, Messenger LA. The effect of next-generation, dual-active-ingredient, long-lasting insecticidal net deployment on insecticide resistance in malaria vectors in Benin: results of a 3-year, three-arm, cluster-randomised, controlled trial. Lancet Planet Health. 2024 Nov;8(11):e894–e905. doi: 10.1016/S2542-5196(24)00232-8. PMID: 39515347.

23. Chen, M., Yan, R., Xu, Z. et al. Knockdown and insecticidal activities of pyrethrum, dimefluthrin and meperfluthrin against Aedes aegypti and Aedes albopictus. Int J Trop Insect Sci 45, 383–397 (2025). 10.1007/s42690-025-01437-3

24. Moyer, B.; Rock, A.N.; Clayton, D.H. Experimental Test of the Importance of Preen Oil in Rock Doves (Columba livia). Auk 2003, 120, 490–496.

25. Williams, W.G.; Kennedy, G.G.; Yamamoto, R.T.; Thacker, J.D.; Bordner, J. 2-Tridecanone: A naturally occurring insecticide from the wild tomato Lycopersicon hirsutum f. glabratum. Science 1980, 207, 888–889.

26. Anuradha, V.; Syed, A.M.; Yogananath, N. Efficacy of Mosquito Repellent and Adulticidal Activities of Halophila ovalis Extract Against Filaria Vectors. J. Trop. Dis. 2016, 4, 1000191.

27. Martínez-de la Puente, Josué & Díez-Fernández, Alazne & Soriguer, Ramón & Rambozzi, Luisa & Peano, Andrea & Meneguz, Pier Giuseppe & Figuerola, Jordi. (2021). Are Malaria-Infected Birds More Attractive to Mosquito Vectors?. Ardeola: International Journal of Ornithology. 68. 205–218. 10.13157/arla.68.1.2021.fo1.

28. Bharadwaj, N., Sharma, R., Subramanian, M., Ragini, G., Nagarajan, S. A., & Rahi, M. (2025). Omics Approaches in Understanding Insecticide Resistance in Mosquito Vectors. International journal of molecular sciences, 26(5), 1854. 10.3390/ijms26051854

29. Avramov, M., Thaivalappil, A., Ludwig, A., Miner, L., Cullingham, C. I., Waddell, L., & Lapen, D. R. (2024). Relationships between water quality and mosquito presence and abundance: a systematic review and meta-analysis. Journal of Medical Entomology, 61(1), 1–33.

30. Fazeli-Dinan, M., Azarnoosh, M., Özgökçe, M. S., Chi, H., Hosseini-Vasoukolaei, N., Haghi, F. M., & Hemingway, J. (2022). Global water quality changes posing threat of increasing infectious diseases, a case study on malaria vector Anopheles stephensi coping with the water pollutants using age-stage, two-sex life table method. Malaria Journal, 21(1), 178.

31. Pathak, A. K., Quek, S., Sharma, R., Shiau, J. C., Thomas, M. B., Hughes, G. L., & Murdock, C. C. (2025). Thermal variation influences the transcriptome of the major malaria vector Anopheles stephensi. Communications Biology, 8(1), 112.

32. Noori, N., Lockaby, B. G., & Kalin, L. (2015). Larval development of Culex quinquefasciatus in water with low to moderate. Journal of Vector Ecology, 40(2), 208–220.

33. Goselle, O. N., Gyang, D. A., Adara, O. F., Effiong, K. T., Nanvyat, N., Barshepn, Y., … & Mafuyai, H. B. (2018). A comparative study of the growth pattern of the larvae of Culex Linnaeus, 1758 in various water bodies and the implications for environmental control.

34. Muturi, E. J., Mwangangi, J., Shililu, J., Jacob, B. G., Mbogo, C., Githure, J., & Novak, R. J. (2008). Environmental factors associated with the distribution of Anopheles arabiensis and Culex quinquefasciatus in a rice agro-ecosystem in Mwea, Kenya. Journal of Vector Ecology, 33(1), 56–63.

35. Onchuru, T. O., Ajamma, Y. U., Burugu, M., Kaltenpoth, M., Masiga, D., & Villinger, J. (2016). Chemical parameters and bacterial communities associated with larval habitats of Anopheles, Culex and Aedes mosquitoes (Diptera: Culicidae) in western Kenya. International Journal of Tropical Insect Science, 36(3), 146–160.

36. Talipouo, A., Mavridis, K., Nchoutpouen, E., et al. High insecticide resistance mediated by different mechanisms in *Culex quinquefasciatus* populations from the city of Yaoundé, Cameroon. Sci Rep 11, 7322 (2021). 10.1038/s41598-021-86850-7

37. Yuan H, Shan W, Zhang Y, Yan H, Li Y, et al. (2023) High frequency of Voltage-gated sodium channel (VGSC) gene mutations in Aedes albopictus (Diptera: Culicidae) suggest rapid insecticide resistance evolution in Shanghai, China. PLOS Neglected Tropical Diseases 17(6): e0011399. 10.1371/journal.pntd.0011399

38. Williams, J., Cowlishaw, R., Sanou, A., Ranson, H. and Grigoraki, L. (2022), In vivo functional validation of the V402L voltage gated sodium channel mutation in the malaria vector An. gambiae. Pest Manag Sci, 78: 1155–1163. 10.1002/ps.6731

39. Huang, Y., Guo, Q., Sun, X., Zhang, C., Xu, N., Xu, Y., Zhou, D., Sun, Y., Ma, L., Zhu, C., & Shen, B. (2018). Culex pipiens pallens cuticular protein CPLCG5 participates in pyrethroid resistance by forming a rigid matrix. Parasites & vectors, 11(1), 6. 10.1186/s13071-017-2567-9

40. Cooperband, M. F., & Allan, S. A. (2009). Effects of different pyrethroids on landing behavior of female Aedes aegypti, Anopheles quadrimaculatus, and Culex quinquefasciatus mosquitoes (Diptera: Culicidae). Journal of medical entomology, 46(2), 292–306.

41. Kauffman, E., Payne, A., Franke, M. A., Schmid, M. A., Harris, E., & Kramer, L. D. (2017). Rearing of Culex spp. and Aedes spp. mosquitoes. Bio-protocol, 7(17), e2542–e2542.

42. CDC: Manual for Evaluating Insecticide Resistance Using the CDC Bottle Bioassay. CDC; 2023.

43. Agumba, S., Gimnig, J. E., Ogonda, L., Ombok, M., Kosgei, J., Munga, S., Guyah, B., Omondi, S., & Ochomo, E. (2019). Diagnostic dose determination and efficacy of chlorfenapyr and clothianidin insecticides against Anopheles malaria vector populations of western Kenya. Malaria journal, 18(1), 243. 10.1186/s12936-019-2858-z

44. Praulins, G., Murphy-Fegan, A., Gillespie, J., Mechan, F., Gleave, K., & Lees, R. (2024). Unpacking WHO and CDC Bottle Bioassay Methods: A Comprehensive Literature Review and Protocol Analysis Revealing Key Outcome Predictors. Gates open research, 8, 56. 10.12688/gatesopenres.15433.1

45. Horikoshi M, Tang Y (2018). ggfortify: Data Visualization Tools for Statistical Analysis Results. https://CRAN.R-project.org/package=ggfortify

46. Lüdecke, D., Ben-Shachar, M. S., Patil, I., Waggoner, P., & Makowski, D. (2021). performance: An R package for assessment, comparison and testing of statistical models. Journal of open source software, 6(60)

47. R Core Team (2024) R: A Language and Environment for Statistical Computing. R Foundation for Statistical Computing.

48. Wickham, H., Chang, W., & Wickham, M. H. (2016). Package ‘ggplot2’. Create elegant data visualisations using the grammar of graphics. Version, 2(1), 1–189.

49. Global vector control response 2017–2030. Geneva: World Health Organization; 2017. Licence: CC BY-NC-SA 3.0 IGO.

50. Kasai, S., Itokawa, K., Uemura, N., Takaoka, A., Furutani, S., Maekawa, Y., Kobayashi, D., Imanishi-Kobayashi, N., Amoa-Bosompem, M., Murota, K., Higa, Y., Kawada, H., Minakawa, N., Cuong, T. C., Yen, N. T., Phong, T. V., Keo, S., Kang, K., Miura, K., Ng, L. C., … Komagata, O. (2022). Discovery of super-insecticide-resistant dengue mosquitoes in Asia: Threats of concomitant knockdown resistance mutations. Science advances, 8(51), eabq7345. 10.1126/sciadv.abq7345

51. World health statistics 2025: monitoring health for the SDGs, Sustainable Development Goals. Geneva: World Health Organization; 2025. Licence: CC BY-NC-SA 3.0 IGO.

## REFERENCES

[1] Franklinos LHV, Jones KE, Redding DW, Abubakar I. The effect of global change on mosquito-borne disease. Lancet Infect Dis 2019;19:e302–e312. 10.1016/S1473-3099(19)30161-6

[2] Teeluck M, Adegboye O, Karl S, Iyaloo DP, McBryde E. Understanding the effect of a changing climate on the re-emergence of mosquito-borne diseases in vulnerable small island nations: a systematic review. Zoonoses Public Health 2025;72:223–247. 10.1111/zph.13212

[3] World Health Organization. Global vector control response 2017–2030. Geneva: World Health Organization; 2017. ISBN 978-92-4-151297-8.

[4] World Health Organization. World malaria report 2025: addressing the threat of antimalarial drug resistance. Geneva: World Health Organization; 2025. ISBN 978-92-4-011782-2

[5] World Health Organization. WHO guidelines for malaria, 10 September 2026. Geneva: World Health Organization; 2026. 10.2471/B09879

[6] Suh PF, Elanga-Ndille E, Tchouakui M, Sandeu MM, Tagne D, Wondji C, Ndo C. Impact of insecticide resistance on malaria vector competence: a literature review. Malar J 2023;22:19. 10.1186/s12936-023-04444-2

[7] Oruni A, Tchouakui M, Djoko Tagne CS, Hearn J, Kayondo J, Wondji CS. Temporal evolution of insecticide resistance and bionomics in *Anopheles funestus*, a key malaria vector in Uganda. Sci Rep 2024;14:32027. 10.1038/s41598-024-83689-6

[8] Teshome A, Erko B, Golassa L, Yohannes G, Irish SR, Zohdy S, Yoshimizu M, Dugassa S. Resistance of *Anopheles stephensi* to selected insecticides used for indoor residual spraying and long-lasting insecticidal nets in Ethiopia. Malar J 2023;22:218. 10.1186/s12936-023-04649-5

[9] Abbasi E, Daliri S, Talbalaghi A, Mehrpouya F, Hasanzadeh Arab M, Aslvaeli A, Moemenbellah-Fard MD. Knockdown resistance (kdr)-associated organochlorine resistance in mosquito-borne diseases (*Culex pipiens*): a systematic review and meta-analysis. Heliyon 2025;11:e41571. 10.1016/j.heliyon.2024.e41571

[10] Soltanbeiglu S, Mohammadi Bavani M, Vahedi M, Chavshin AR. Susceptibility of *Culiseta longiareolata* to insecticides in West Azerbaijan province, northwestern Iran. Vet Res Forum 2022;13:435–437. 10.30466/vrf.2020.134833.302

[11] Asgarian TS, Vatandoost H, Hanafi-Bojd AA, Nikpoor F. Worldwide status of insecticide resistance of *Aedes aegypti* and *Aedes albopictus*, vectors of arboviruses of chikungunya, dengue, Zika and yellow fever. J Arthropod Borne Dis 2023;17:1–27. 10.18502/jad.v17i1.13198

[12] Mora-Rubio C, García-Longoria L, Marzal A. Preening for protection: a systematic review of the antimicrobial properties of uropygial secretions. J Avian Biol 2025;2025:e03425. 10.1111/jav.03425

[13] Magallanes S, Møller AP, García-Longoria L, de Lope F, Marzal A. Volume and antimicrobial activity of secretions of the uropygial gland are correlated with malaria infection in house sparrows. Parasit Vectors 2016;9:232. 10.1186/s13071-016-1512-7

[14] Marzal A, Møller AP, Espinoza K, Morales S, Luján-Vega C, Cárdenas-Callirgos JM, Mendo L, Álvarez-Barrientos A, González-Blázquez M, García-Longoria L, et al. Variation in malaria infection and immune defence in invasive and endemic house sparrows. Anim Conserv 2018;21:505–514. 10.1111/acv.124

[15] Magallanes S, Møller AP, Luján-Vega C, Fong E, Vecco D, Flores-Saavedra W, García-Longoria L, de Lope F, Iannacone JA, Marzal A. Exploring the adjustment to parasite pressure hypothesis: differences in uropygial gland volume and haemosporidian infection in Palearctic and Neotropical birds. Curr Zool 2021;67:147–156. 10.1093/cz/zoaa037

[16] Marzal A, Magallanes S, García-Longoria L. Stimuli followed by avian malaria vectors in host-seeking behaviour. Biology (Basel) 2022;11:726. 10.3390/biology11050726

[17] Díez-Fernández A, Martínez-de la Puente J, Gangoso L, Ferraguti M, Soriguer R, Figuerola J. House sparrow uropygial gland secretions do not attract ornithophilic nor mammophilic mosquitoes. Med Vet Entomol 2020;34:225–228. 10.1111/mve.12401

[18] Cai P, Dimopoulos G. Microbial biopesticides: a One Health perspective on benefits and risks. One Health 2025;20:100962. 10.1016/j.onehlt.2024.100

[19] Kauffman E, Payne A, Franke MA, Schmid MA, Harris E, Kramer LD. Rearing of *Culex* spp. and *Aedes* spp. mosquitoes. Bio Protoc 2017;7:e2542. 10.21769/BioProtoc.2542

[20] Centers for Disease Control and Prevention. Manual for evaluating insecticide resistance using the CDC bottle bioassay. Atlanta, GA: Centers for Disease Control and Prevention; 2012.

[21] Praulins G, et al. Unpacking WHO and CDC bottle bioassay methods: a comprehensive literature review and protocol analysis revealing key outcome predictors. Gates Open Res 2024;8:56. 10.12688/gatesopenres.15433.2

[22] Tang Y, Horikoshi M, Li W. ggfortify: unified interface to visualize statistical result of popular R packages. R J 2016;8:474–485. 10.32614/RJ-2016-06

[23] Lüdecke D, Ben-Shachar MS, Patil I, Waggoner P, Makowski D. performance: an R package for assessment, comparison and testing of statistical models. J Open Source Softw 2021;6:3139. 10.21105/joss.03139

[24] R Core Team. R: a language and environment for statistical computing. Vienna: R Foundation for Statistical Computing; 2024

[25] Wickham H, Chang W, Wickham M. Package “ggplot2”. Create elegant data visualisations using the grammar of graphics (Versión 2). r-project.org. 2016

[26] Sovi A, Adoha CJ, Yovogan B, Cross CL, Dee DP, Konkon AK, Sidick A, Accrombessi M, Ahouandjinou MJ, Ossè R, et al. The effect of next-generation, dual-active-ingredient, long-lasting insecticidal net deployment on insecticide resistance in malaria vectors in Benin: results of a 3-year, three-arm, cluster-randomised, controlled trial. Lancet Planet Health 2024;8:e894–e905. 10.1016/S2542-5196(24)00232

[27] Chen M, Yan R, Xu Z, Zhou Q, et al. Knockdown and insecticidal activities of pyrethrum, dimefluthrin and meperfluthrin against *Aedes aegypti* and *Aedes albopictus*. Int J Trop Insect Sci 2025;45:383–397. 10.1007/s42690-025-01437-3

[28] Moyer BR, Rock AN, Clayton DH. Experimental test of the importance of preen oil in rock doves (*Columba livia*). Auk 2003;120:490–496. 10.1093/auk/120.2.490

[29] Williams WG, Kennedy GG, Yamamoto RT, Thacker JD, Bordner J. 2-Tridecanone: a naturally occurring insecticide from the wild tomato *Lycopersicon hirsutum* f. *glabratum*. Science 1980;207:888–889. 10.1126/science.207.4433.888

[30] Anuradha V, Syed AM, Yogananath N. Efficacy of mosquito repellent and adulticidal activities of *Halophila ovalis* extract against filaria vectors. J Trop Dis 2016;4:1000191

[31] Bharadwaj N, Sharma R, Subramanian M, Ragini G, Nagarajan SA, Rahi M. Omics approaches in understanding insecticide resistance in mosquito vectors. Int J Mol Sci 2025;26:1854. 10.3390/ijms26051854

[32] Avramov M, Thaivalappil A, Ludwig A, Miner L, Cullingham CI, Waddell L, Lapen DR. Relationships between water quality and mosquito presence and abundance: a systematic review and meta-analysis. J Med Entomol 2024;61:1–33. 10.1093/jme/tjad139

[33] Fazeli-Dinan M, Azarnoosh M, Özgökçe MS, Chi H, Hosseini-Vasoukolaei N, Motevalli Haghi F, Zazouli MA, Nikookar SH, Dehbandi R, Enayati A, et al. Global water quality changes posing threat of increasing infectious diseases, a case study on malaria vector *Anopheles stephensi* coping with water pollutants using age-stage, two-sex life table method. Malar J 2022;21:178. 10.1186/s12936-022-04201-x

[34] Pathak AK, Quek S, Sharma R, Shiau JC, Thomas MB, Hughes GL, et al. Thermal variation influences the transcriptome of the major malaria vector *Anopheles stephensi*. Commun Biol 2025;8:112. 10.1038/s42003-025-07477-2

[35] Becker N, Petrić D, Zgomba M, Boase C, Madon MB, Dahl C, Kaiser A. Mosquitoes: identification, ecology and control. 3rd ed. Cham: Springer; 2020. 10.1007/978-3-030-11623-1

[36] Onchuru TO, Ajamma YU, Burugu M, Kaltenpoth M, Masiga D, Villinger J. Chemical parameters and bacterial communities associated with larval habitats of *Anopheles*, *Culex* and *Aedes* mosquitoes (Diptera: Culicidae) in western Kenya. Int J Trop Insect Sci 2016;36:146–160

[37] Talipouo A, Mavridis K, Nchoutpouen E, Djiappi-Tchamen B, Fotakis EA, Kopya E, Bamou R, Kekeunou S, Awono-Ambene P, Balabanidou V, et al. High insecticide resistance mediated by different mechanisms in Culex quinquefasciatus populations from the city of Yaoundé, Cameroon. Sci Rep 2021;11:7322. 10.1038/s41598-021-86850-7

[38] Yuan H, Shan W, Zhang Y, Yan H, Li Y, Zhou Q, Dong H, Tao F, Liu H, Leng P, et al. High frequency of voltage-gated sodium channel (VGSC) gene mutations in *Aedes albopictus* suggests rapid insecticide resistance evolution in Shanghai, China. PLoS Negl Trop Dis 2023;17:e0011399. 10.1371/journal.pntd.0011399

[39] Williams J, Cowlishaw R, Sanou A, Ranson H, Grigoraki L. In vivo functional validation of the V402L voltage-gated sodium channel mutation in the malaria vector *Anopheles gambiae*. Pest Manag Sci 2022;78:1155– 1163. 10.1002/ps.673

[40] Huang Y, Guo Q, Sun X, Zhang C, Xu N, Xu Y, Zhou D, Sun Y, Ma L, Zhu C, Shen B. *Culex pipiens pallens* cuticular protein CPLCG5 participates in pyrethroid resistance by forming a rigid matrix. Parasit Vectors 2018;11:6. 10.1186/s13071-017-2567-9

[41] Cooperband MF, Allan SA. Effects of different pyrethroids on landing behavior of female *Aedes aegypti*, *Anopheles quadrimaculatus* and *Culex quinquefasciatus* mosquitoes (Diptera: Culicidae). J Med Entomol 2009;46:292–306. 10.1603/033.046.0214

